# Modeling Post-traumatic memory deficits: Repeated head injury effects on Novelty detection in *Drosophila*

**DOI:** 10.64898/2026.04.30.721976

**Authors:** Prachi Shah, Aparna Dev, Brittany Lew, Christina Biuckians, Anne S. Oepen, Alessia Ghirelli, Tamara Boto, Isaac Cervantes-Sandoval

## Abstract

Traumatic brain injury (TBI) is a leading cause of neurological dysfunction, yet the mechanisms linking repeated concussions to cognitive impairment remain poorly defined. Here, we used a controlled *Drosophila* model of repetitive head trauma using a piezoelectric actuator to deliver reproducible mild, moderate, or severe concussions. Repeated trauma significantly reduced lifespan and induced transient locomotor deficits, persistent motivational impairments, and surprisingly sleep behavior was only altered with injury protocols expanding across more than one day. At the cellular level, concussions triggered a delayed but robust proliferation of astrocyte-like glia, consistent with neuroinflammatory responses observed in mammalian models. To investigate circuit-level consequences, we examined novelty detection within the mushroom body, focusing on MBON-α3, a neuron that shows plasticity consistent with odor habituation similar to previously reported for MBON-α’3. Functional calcium imaging revealed that concussed flies exhibited disrupted odor-induced plasticity and diminished baseline responsiveness in MBON-α3 at one- and five- days post-injury, despite intact Kenyon cell input. Notably, these deficits were force-dependent and largely reversible by ten days, highlighting both vulnerability and resilience within this defined circuit. Together, our findings demonstrate that repeated concussions in *Drosophila* produce dose-dependent behavioral, cellular, and circuit dysfunctions that parallel mammalian TBI. This work establishes a genetically tractable platform for dissecting the mechanisms of concussion-induced cognitive decline and for identifying potential targets for intervention.

## Introduction

Traumatic brain injury (TBI) is a significant public health concern and a major cause of long-term neurological disability worldwide (Dewan et al., 2018). Among its many forms, repetitive mild to moderate head trauma, often referred to as repeated concussions, has garnered increasing attention due to its association with chronic neurocognitive impairments and neurodegenerative diseases such as chronic traumatic encephalopathy (CTE) (McKee et al., 2013). Despite its clinical relevance, the mechanistic understanding of how repetitive head trauma affects brain cognitive functions remains incomplete, largely due to the heterogeneity of the injure nature and the complexity and limitations of current mammalian models. As such, *Drosophila melanogaster* has emerged as a powerful, genetically tractable model organism to study the cellular and behavioral consequences of brain injury (Katzenberger et al., 2013; McKee et al., 2013; Barekat et al., 2016; Saikumar et al., 2020).

*Drosophila* offers numerous advantages for modeling TBI, which have been recently reviewed by Kelly et al. (Kelly et al., 2025): a well-characterized nervous system, sophisticated genetic tools, short generation time, and the availability of high-throughput behavioral and imaging assays (Duffy, 2002). Importantly, fly models of TBI have demonstrated parallels with mammalian injury responses, including neuroinflammation, glial activation, behavioral deficits, and reduced lifespan (Katzenberger et al., 2013; van Alphen et al., 2022). These features make *Drosophila* an ideal model to dissect the cellular mechanisms and functional outcomes of head trauma.

In recent years, novel methodologies have enabled researchers to apply controlled head-specific concussions to *Drosophila*. In particular, Saikumar et al., (Saikumar et al., 2020, 2021) developed a head-targeted TBI protocol using a piezoelectric actuator, allowing precise and repeatable concussive forces to be delivered directly to the fly head while minimizing systemic injuries. This advancement allows the modeling of graded head trauma, mild, moderate, or severe, and the evaluation of dose-dependent effects of TBI. In parallel, Hattori et al. (Hattori et al., 2017) introduced a functional imaging approach using calcium indicators to monitor odor-evoked responses in identified mushroom body output neurons (MBONs), specifically MBON-α’3, which is known to be involved in olfactory novelty detection and memory formation. These two methods together provide a powerful framework for investigating how repetitive head injury affects a well-defined simple neural circuit that function in a cognitive task such as novelty detection.

In this study, we employed these cutting-edge approaches to examine the impact of repeated head concussions on survival, locomotor behavior, mood-related behaviors, glial activation, and cognitive function in *Drosophila melanogaster*. We found that flies subjected to mild and moderate concussions exhibited reduced neuronal plasticity at both 1- and 5-days post-injury but showed complete recovery by day 10. Importantly, these deficits cannot be explained by impaired sensory perception, as Kenyon cell responses remained intact. Instead, our results suggest that repeated head concussions primarily disrupt the ability of flies to induce synaptic plasticity within a defined neural circuit.

Together, these results demonstrate that repeated head trauma in *Drosophila* causes a spectrum of dose-dependent physiological, behavioral, and cognitive impairments, paralleling the effects observed in mammalian models and human patients. Importantly, our findings establish olfactory novelty detection and MBON-α3 activity as sensitive and quantifiable readouts of post-concussive cognitive dysfunction. Furthermore, the robust activation and proliferation of astrocyte-like glia following injury support the utility of *Drosophila* in modeling neuroinflammatory contributions to TBI-related cognitive decline.

In summary, our study contributes to a growing body of work positioning *Drosophila melanogaster* as a powerful in vivo platform for studying the cellular and circuit-level consequences of traumatic brain injury, with the added benefit of precise genetic and functional imaging tools. These results provide valuable insights into the mechanisms underlying TBI and lay the groundwork for future genetic dissection of resilience and vulnerability to concussion-induced brain dysfunction.

## Results and Discussion

Traumatic brain injury has been amply studied in the context of immune system activation, inflammation, neuronal degeneration, and its long-term behavioral consequences (McKee et al., 2013). Less is understood about how the initial stages post trauma affects cognitive processes in well-defined neuronal circuits. Here, we decided to explore the effects of repeated head concussion on well-defined memory circuit involved in cognitive task of novelty odor detection in the fruit fly brain. Even though, the complete neuronal circuit computing novelty detection is more complex, this can be simplistically reduced to the main components which is composed of three-layer circuit. In principle, odors are first represented by sparse coding in a reduced number of KC (∼10%) (Honegger et al., 2011). MB convey this activity to a handful number of MBON that tile the KC axonal projections (Aso et al., 2014a, 2014b). These KC>MBON tiles receive inputs from dopaminergic neurons that are arranged in an almost mirror-like pattern, forming ∼15 defined compartments (Mao and Davis, 2009; Aso et al., 2014a, 2014b). It is thought that each of these compartments is subjected to experience driven plasticity and the combinatorial integration of the overall output of these compartments defined the expression of learn behavior (Séjourné et al., 2011; Pai et al., 2013; Plaçais et al., 2013; Bouzaiane et al., 2015; Hige et al., 2015b, 2015a; Owald et al., 2015; Berry et al., 2018; Cervantes-sandoval et al., 2020; Martinez-Cervantes et al., 2022). At least two of these compartments but possible more, MBON-α’3 (Hattori et al., 2017) and MBON-α3 (This study) can be implicated in detection of olfactory novelty. When flies encounter a novel odor, MBON-α’3 respond strongly, but this activity diminishes rapidly with repeated presentations of the same odor. This shift in neural responsiveness during familiarization depends on odor-driven activity of the counterpart dopaminergic neuron that innervates the same compartment. Hattori et al. (2017) also reported that novel odors also trigger an alerting behavioral reaction, which disappears when these MBONs are silenced. These findings indicate that the α’3 compartment contributes causally to how flies distinguish novel from familiar odor stimuli, through dopamine-dependent plasticity at the KC–MBONα‘3 synapse. Here we tested the effects of repeated head trauma in this simple three layer circuit encoding novelty detection. First, we built and validated a concussion apparatus based on (Saikumar et al., 2020, 2021) Briefly, a small piezo actuator is placed over flies immobilized in a collar. An Increasing amount of current is applied using an Arduino microcontroller to deflect downwards the piezo element in a force and temporally controlled manner (Figure 1A). This allows a precise and reproducible fly head compression. Using this device, we concussed flies with three different forces (mild-35%, moderate-40%, and severe-45%), five times with one min ISI. Experiments included a sham control, where the flies were immobilized in the collar for the same time the concussion takes but not head compression was delivered. This sham control accounts for introducing the flies to the collar. One, five or ten days after head concussions, different parameters were measured, including survival curves, climbing assay, glia proliferation, sleep, force swim test and using functional imaging, olfactory novelty detection (Figure 1B). Repeated head concussions led to a marked reduction in lifespan across all levels of force when compared with sham-treated control flies, as determined by both the log-rank and Gehan–Breslow–Wilcoxon survival tests (Figure 1C). Median survival declined substantially with injury severity: flies subjected to mild repeated concussions survived a median of 34 days, compared with 59 days for matched sham controls; those exposed to moderate concussions survived only 13 days versus 63 days for controls; and flies experiencing severe concussions had a median survival of 29.5 days compared with 60 days in their respective controls. Notably, much of the divergence between survival curves arose from an acute wave of mortality occurring shortly after the concussion event. Beyond this early phase, survival rates appeared to stabilize, with the rate of decline partially normalizing approximately 20 days post-injury (Figure 1C). These results indicate that repeated head concussions were reliable in inducing premature mortality at all forces tested. Based on these findings, we focused most subsequent experiments on the two extreme force conditions, mild and severe, except in the case of functional imaging, where all force levels were included.

**Figure 1.**
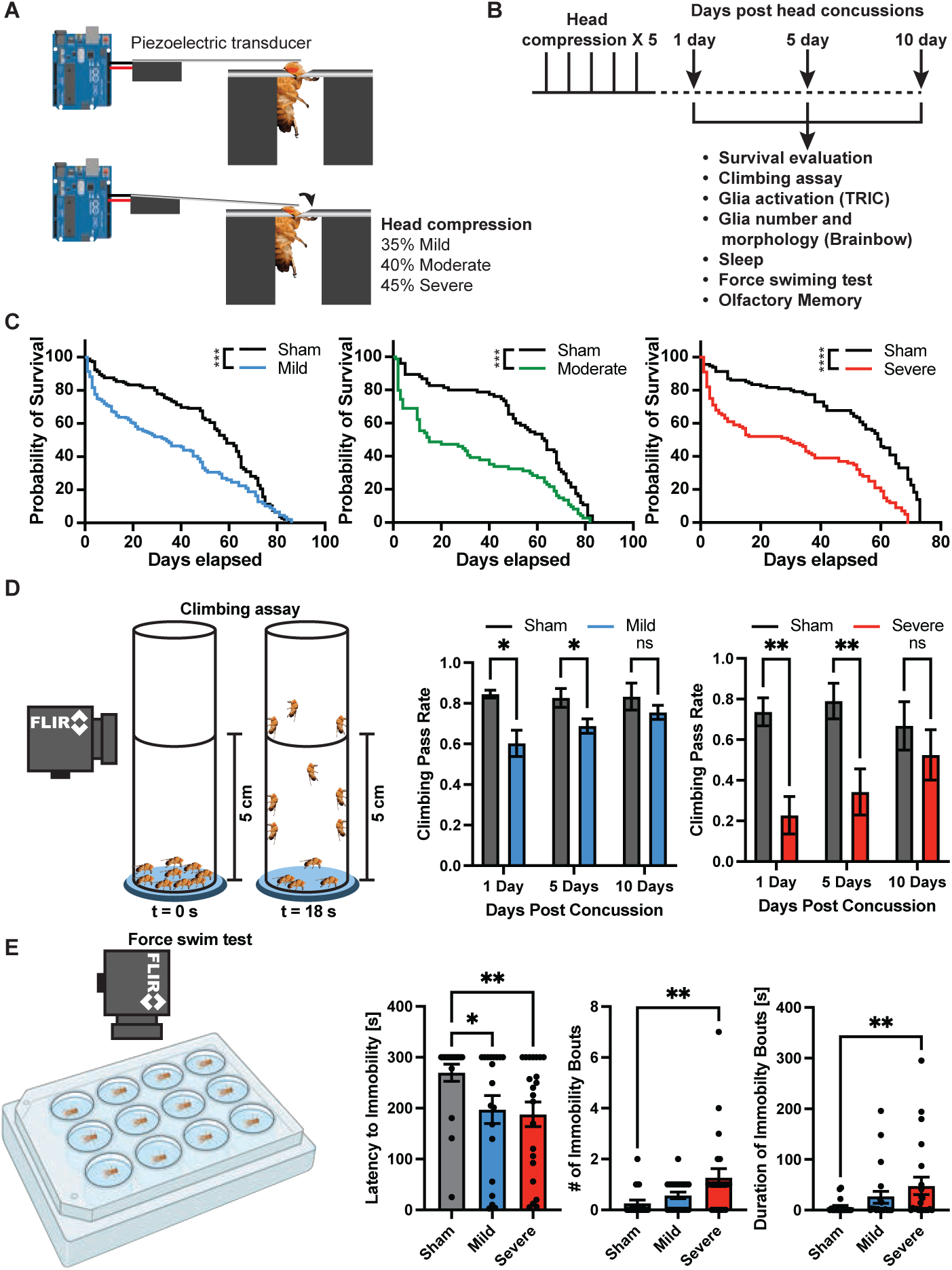
Behavioral effects to repeated controlled head trauma. (**A**) Schematic of a piezoelectric transducer–based head trauma device operated via an Arduino microcontroller. (**B**) Experimental design. Flies were repeatedly concussed five times every minute and their survival tracked. One, five- or ten-days post head trauma flies were evaluated for climbing, glia activation and proliferation, sleep, force swim test, and olfactory memory. (**C**) Survival curves after different severity of head concussions compared to match sham flies. Mild, moderate and severe concussions were significantly different compared to sham flies. Survival Log-rank (Mantel-Cox) test, \**p<0.0001.* (**D**) Left, diagram of climbing assay. Right, flies were evaluated for negative geotaxis one-, five- and ten-days post head trauma. Both mild and severe concussions have significant impaired locomotion, one- and five-days post head trauma when compared to match sham flies. Flies seem to recover with time as climbing capacity recovers ten-days post head trauma. Two-way ANOVA, followed by Fisher multiple comparison, *\*p<0.05, **p<0.004.* (**E**) Left, diagram of forced swim test. Right, Flies showed significantly impaired latency to immobility following mild and severe repeated head trauma. Increased number and duration of immobility bouts was observed only severe repeated head trauma. One-way ANOVA, Kluskal-Wallis test followed by Dunn’s multiple comparison, *\*p<0.05, **p<0.008*.

Using the climbing assay to assess motor performance (Figure 1D), we observed that flies subjected to repeated mild concussions displayed a modest but significant reduction in climbing success one and five days after injury. Interestingly, by ten days post-concussion, these flies had fully regained their climbing ability, indicating a transient motor deficit (Figure 1D, middle panel). In contrast, flies exposed to repeated severe concussions showed a much more pronounced impairment in climbing capacity at both one and five days after injury. Surprisingly, however, even these severely injured flies exhibited a full recovery of negative geotaxis by ten days post-concussion, suggesting that despite the acute and force-dependent motor deficits, flies can regain locomotor function over time. These results suggest that while repeated concussions cause lasting reductions in survival, their effects on motor performance are largely reversible.

To assess whether head concussions affected motivational state beyond their impact on locomotor ability, we employed the forced swim test (FST) (Hibicke and Nichols, 2022) ten days after injury (Figure 1E). In this assay, individual flies were gently aspirated into wells of a 24-well plate filled with water and their behavior recorded using a FLIR camera. Three behavioral parameters were quantified: latency to immobility, number of immobility bouts, and duration of immobility bouts. We found that both mildly and severely concussed flies exhibited a significant reduction in latency to immobility, indicating a faster onset of inactivity compared with sham controls. However, only flies subjected to severe concussions showed significant increases in the number and duration of immobility bouts, whereas flies with mild concussions were indistinguishable from controls on these measures (Figure 1E). These results suggest that while both mild and severe concussions lead to changes in motivational state, the effects are more persistent and pronounced following severe injury. Moreover, even though concussed flies eventually regained normal locomotor capacity in the negative geotaxis assay, the FST revealed that severe concussions leave a lingering deficit in motivational state.

We next examined whether repeated head concussions affected sleep patterns (Figure 2). Surprisingly, flies subjected to either mild or severe concussions with our protocol showed no significant changes in total sleep duration, or in day- and night-time sleep, when compared to sham and naïve controls. Additional sleep parameters, including the number of sleep bouts, average bout length, and day- or night-time activity, also showed no significant differences (data not shown). As sleep disturbances are a common symptom in patients and a reported outcome of head concussions in *Drosophila* and other animal models (Barekat et al., 2016; van Alphen et al., 2022; Morris et al., 2024), we proceeded to induce repetitive mild TBI during two consecutive days (Figure 2D). This recurrent injury paradigm led to an overall reduction of sleep (Figure 2E-F). The effect was stronger during the light phase, were sleep architecture was also affected (Figure 2H-I): flies subjected to repetitive mild TBI for two days displayed more interrupted day sleep, with a trend to increase the number of sleep bouts and a significant reduction on the average sleep bout duration.

**Figure 2.**
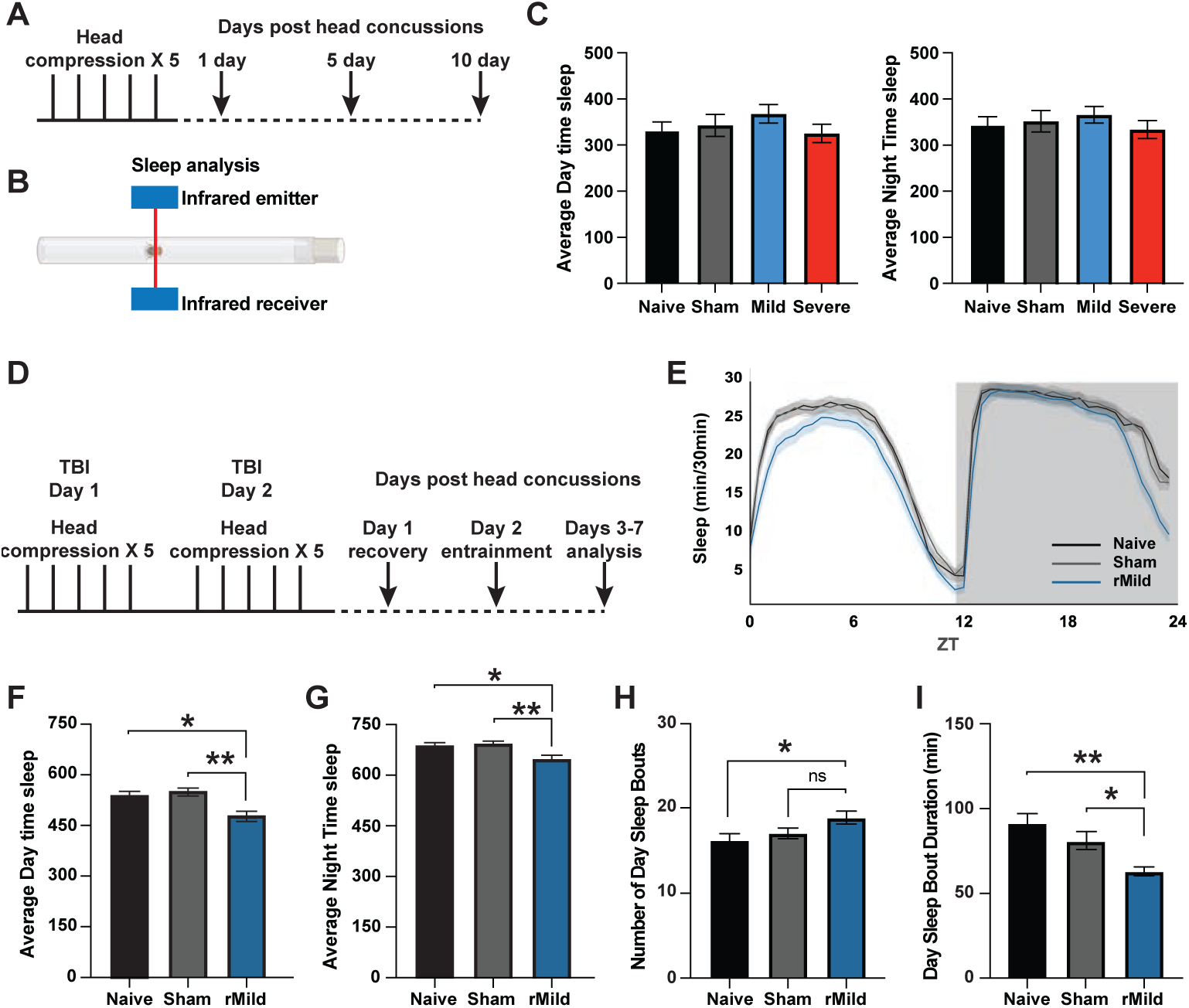
Sleep alterations following repeated head trauma. (**A,B**) Experimental design for 1-day head trauma induction. (**C**) Flies subjected to 1 day of either mild or severe concussions showed no significant changes in total sleep duration, in day- and night-time sleep, when compared to sham and naïve controls. (**D**) Experimental design for 2-day head trauma induction. (**E**) Total sleep in 30 min bins for naïve, sham and repetitive mild TBI (rmTBI) across 5 consecutive days. (**F,G**) Flies subjected to 2 days of mild concussions showed a significant decrease in both day and night-sleep duration when compared to sham and naïve controls. (**H**) Average number of sleep bouts ring the day and (**I**) average day sleep bout duration point towards a more fragmented day sleep in flies following rmTBI. One-way ANOVA followed by Tukey’s multiple comparisons *\*p < 0.05*, *\*\*p < 0.01*, and *\*\*\*p < 0.001*.

Given that repeated concussions produced lasting reductions in survival and persistent changes in motivational state, we next sought to examine their cellular consequences, focusing on glial responses. To approximate changes in glial proliferation following head trauma, we employed the multicolor FlipOut (MCFO) system (Nern et al., 2015) to label astrocyte-like glia in the *Drosophila* brain across different injury severities. This method allowed for stochastic labeling of individual glial cells using the *r86e01-gal4* driver (Figure 3A) (Nern et al., 2011; Kremer et al., 2017). Briefly, flies were subjected to a 10-minute heat shock at 36 °C during the late pupal stage to induce sparse labeling. After eclosion, adult flies were exposed to repeated concussions of varying intensities, and glial cell number was assessed at one, five-, and ten-days post-injury. To facilitate accurate quantification, astroglia were stained with two different antibodies, allowing visualization of individual cells in up to three distinct colors. This multicolor labeling strategy ensured that neighboring cells could be easily distinguished, greatly improving precision in cell counts.

**Figure 3.**
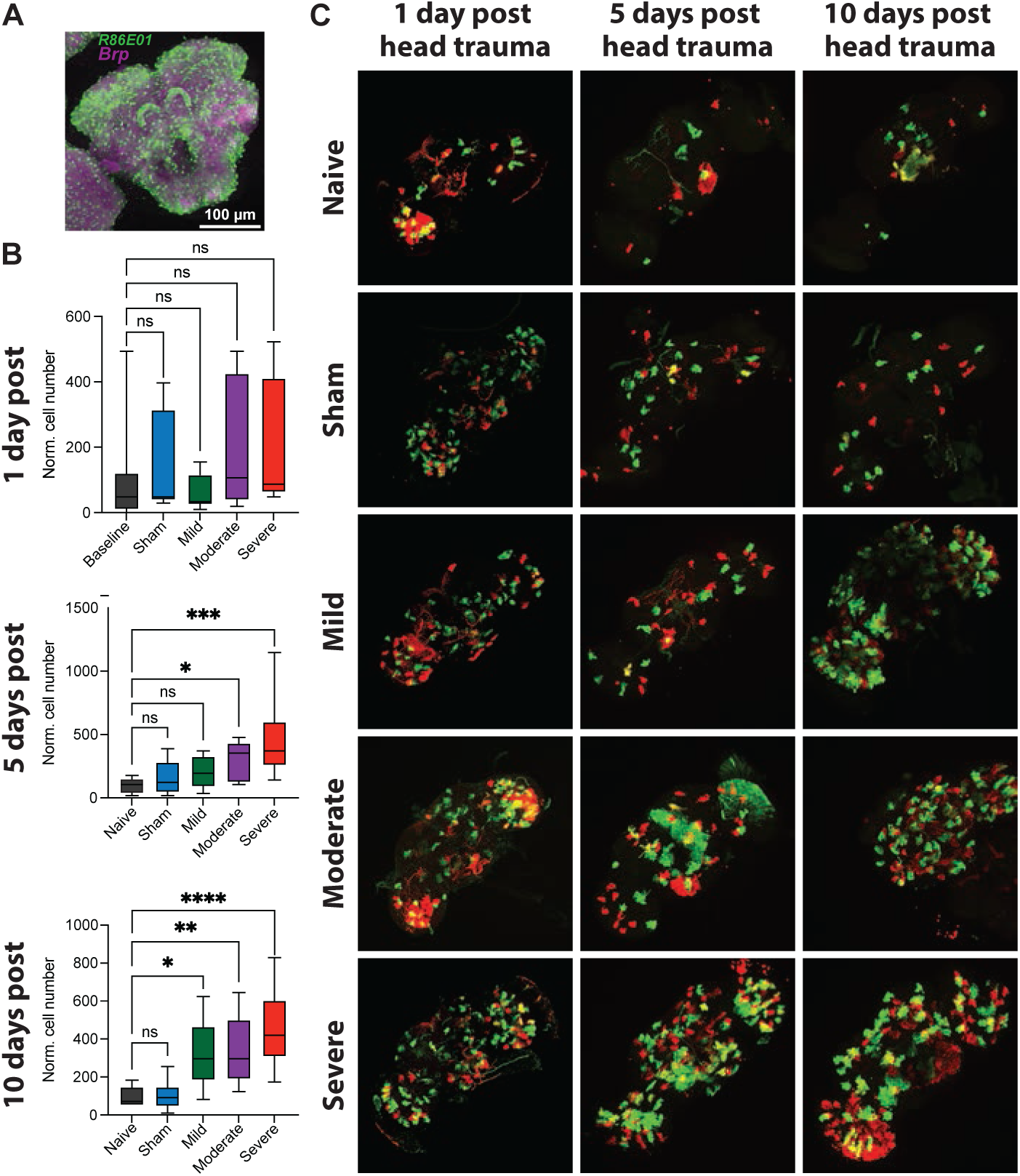
Glia proliferation following repeated head trauma. (**A**) Immunohistochemistry of expression pattern of gal4 line used to label astrocyte-like glia in the *Drosophila* brain. (**B**) Flies showed no increase in astrocyte-like glia cell number one day after repeated head concussions. Conversely, flies showed a significant increase in astrocyte-like glia cell number five days after repeated head concussions with moderate and severe intensities. Finally, flies showed a significant increase in astrocyte-like glia cell number ten days after repeated head concussions with mild, moderate and severe intensities. One-way Anova, Kluskal-Wallis test followed by Dunn’s multiple comparison, *\*p<0.01, **p<0.0062, ****p<0.0001.* (**C**) Representative images of glia labeled with multicolor flip-out approach.

At one day after concussion, astroglial cell numbers did not differ significantly among flies exposed to mild, moderate, or severe concussions, nor from sham-treated or naïve controls (Figure 3B–C). However, by five days post-injury, flies subjected to moderate and severe concussions exhibited a clear increase in astroglial cell number relative to naïve flies, while mildly concussed and sham-treated flies remained comparable to baseline. By ten days post-concussion, this effect had become more pronounced: all injury groups, including mild concussions, showed a significant elevation in astroglial counts compared with both naïve and sham controls. Together, these results demonstrate that repeated head trauma in *Drosophila* induces a time-dependent increase in astroglial proliferation. The early absence of changes followed by robust increases at later timepoints suggests that glial activation is not an immediate but rather is response that emerges days after trauma. These findings complement our behavioral analyses by showing that, although flies can recover certain functions such as locomotor performance, concussions also trigger progressive cellular changes in the brain. This delayed and persistent activation of glia points to a neuroinflammatory response that may underlie the long-term consequences of repeated head trauma.

After validating our repeated head trauma model in *Drosophila* through behavioral and cellular analyses, we next examined its impact on a neuronal circuit involved in memory and novelty detection. Specifically, we focused on mushroom body output neuron (MBON)-α3, which here, we found it shows similarities to the better characterized MBON-α’3 circuit implicated in encoding novelty responses (Hattori et al., 2017). To do this, we concussed flies of varying severities and performed functional calcium imaging to measure MBON-α3 responses to repeated odor presentation (Figure 3A).

Previous work by Hattori et al. (2017) demonstrated that repeated odor presentation with a 12 s inter-stimulus interval (ISI) leads to suppression of calcium responses in MBON-α’3, but not in other MBON types such as MBON-γ1pedc>α/μ, MBON-μ’2mp, or MBON-μ2μ’2a. Here, we found for the first time that in naïve flies MBON-α3 exhibited a significant decrease in calcium responses following repeated odor exposure with a 12 s ISI (Figure 3A–B). Although the depression in the responses is not complete, this neuron partially resembling the plasticity observed in MBON-α’3.

To better quantify this form of plasticity, we compared the average calcium response to the first three odor presentations with the average response to the last three presentations, as previously described by Hattori et al. (2017). As expected, naïve flies showed a robust reduction in MBON-α3 responses across all timepoints examined (1-, 5-, and 10-days post-injury; Figure 3C, Supp. Fig. 1). Sham-treated flies, however, displayed a transient impairment: they failed to show plasticity at one day post-injury, but recovered fully by five and ten days.

In contrast, flies exposed to mild, moderate, or severe concussions exhibited significant impairments in MBON-α3 plasticity at both one- and five-days post-injury. Remarkably, by ten days post-concussion, all injured groups recovered, suggesting that while repeated head trauma disrupts odor-induced plasticity in the short term, flies retain the capacity for recovery over longer timescales.

Beyond changes in plasticity, we also noticed that the overall magnitude of olfactory responses in MBON-α3 appeared diminished in concussed flies, independent of repeated odor exposure (Figure 3 Supplement 1). To directly test this, we compared the average magnitude of the initial three odor-evoked calcium responses across groups. At both one- and five-days post-injury, all treatments—including sham, mild, moderate, and severe concussions—showed significantly reduced responses compared with naïve flies. By ten days, sham and mildly concussed flies recovered their response magnitude, whereas moderate and severely concussed flies continued to exhibit persistently reduced responses (Figure 4A). These results suggest that, in addition to impaired experience-dependent plasticity, concussed flies also suffer from a sustained reduction in baseline olfactory responsiveness, particularly at higher injury severities.

**Figure 4.**
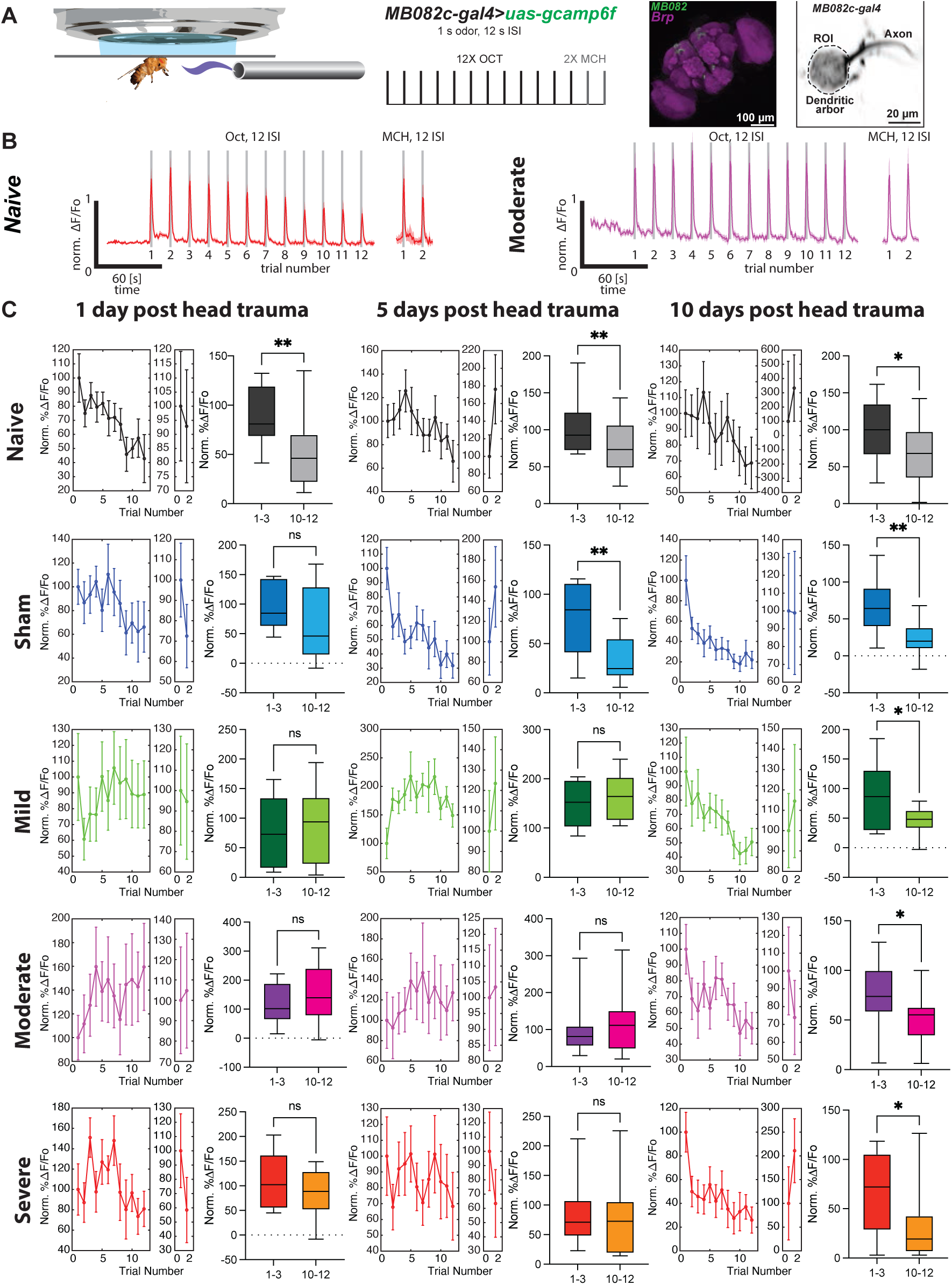
Repeated head trauma impairs neuronal plasticity in a well-defined olfactory circuit. (**A**) Left, diagram of in vivo functional imaging. Middle, experimental protocol. Flies expressing GCaMP6f in MBON-α3 (MB082-gal4) were trained by repeated presentation of 12, 1 s pulses of 1-octanol with a 12 s inter-stimulus interval (ISI), followed by 2 pulses of 4-methyl-cyclohexanol. Right, Immunohistochemistry of expression pattern of MB082-gal4 line used to label MBON-α3 in the *Drosophila* brain and live imaging view of ROI registered during olfactory recordings. (**B**) Functional imaging of normalized calcium traces of MBON-α3 responses upon repeated odor presentations (gray bars) in a naïve animal (red traces) and an animal following moderate repeated head trauma one day after concussions (magenta traces). Shaded area represents SEM. (**C**) Average calcium responses measured in ROI after 12 repeated presentations of OCT and 2 of MCH. Integrated ΔF/Fo signal was normalized for first response. Recordings were obtained in flies one, five- and ten-days post repeated head trauma with different head compressions, sham, mild, moderate and severe. An age-matched naïve control was also used. The error bars represent ± SEM in all figures. Bar graphs show the average of first three responses compared to the last three. As expected, naïve animals showed a significantly decreased responses when first three are compared to last three responses, indicating neuronal plasticity (habituation) after repeated odor presentations from day one to day 10. Interestingly, sham, mild moderate and severe treatments showed impaired neuronal plasticity one day post head trauma. Nevertheless, Sham flies recovered completely five and 10 days after, whereas mild moderate and severe concussed flies remain maintain impairment plasticity 5 days after head concussion. Finally, all flies seem to recover neuronal plasticity 10 days after repeated head trauma as indicated by a significant decrease in odor responses in last three odor presentations compared to first three. Matched-pair T-test, *\*p≤0.0388, **p≤0.0045*.

**Figure 5.**
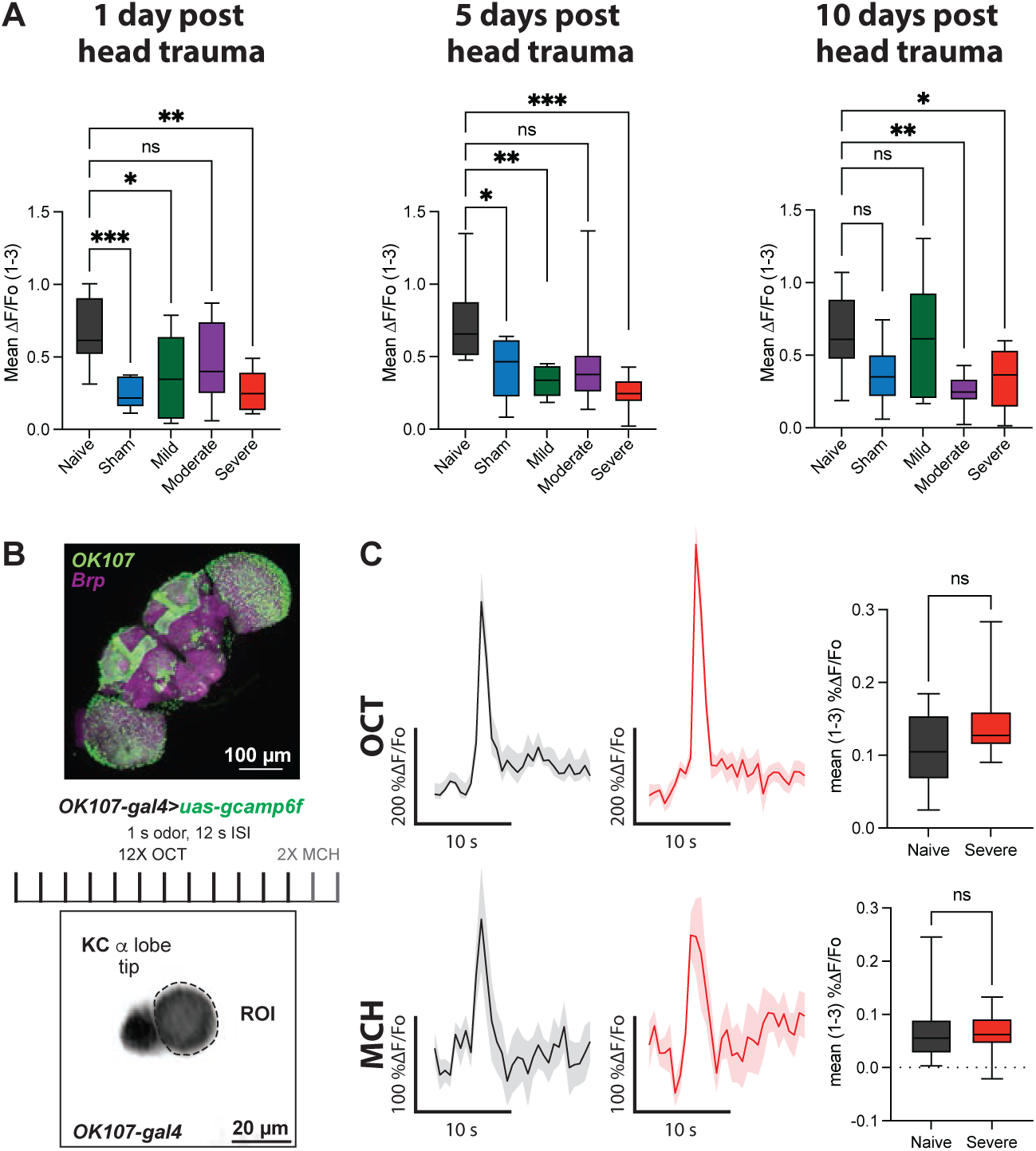
Effect of repeated head trauma on olfactory responses in Kenyon cells. (**A**) Average calcium traces in response to first three odor presentations to OCT and MCH odors. No significant difference in responses between naïve flies and flies subjected to severe head trauma was observed in Kenyon cells. Mann-Whitney test. (**B**) Upper panel, immunohistochemistry of expression pattern of OK107-gal4 line used to label Kenyon cells in the *Drosophila* brain and live imaging view of ROI registered during olfactory recordings in the tip of the α lobe. (**C**) Average response of three odor presentation was significantly lower in MBON-α3 after sham, mild, and severe repeated head trauma one- and five-days post concussions. In contrast, differences were only significant for moderate and severe treatments 10-days after repeated head trauma. One-way ANOVA followed by Dunnett’s multiple comparisons. *\*p<0.014, **p<0.0016, ***p<0.0003*.

One possible explanation for this reduced MBON-α3 activity, is diminished sensory input from Kenyon cells (KCs), the principal intrinsic neurons of the mushroom body. To test this, we recorded olfactory calcium responses directly in KCs at the tip of the α lobe, the region providing input to MBON-α3 (Figure 4B). Surprisingly, even in severely concussed flies, KC responses were indistinguishable from those of naïve controls (Figure 4C). Thus, at least at the KC level, and likely in upstream sensory layers, odor-evoked activity remains intact after head trauma.

Taken together, these findings indicate that repeated concussions impair both odor-induced plasticity and the baseline magnitude of olfactory responses in MBON-α3. Importantly, these deficits cannot be attributed to reduced sensory input from KCs, pointing instead to injury-induced dysfunction at the MBON level or downstream circuit components.

In summary, repeated concussions in *Drosophila* produced consistent and force-dependent effects across multiple levels of analysis: reduced lifespan, transient but reversible locomotor impairments, persistent motivational deficits, increased astroglial proliferation, and disrupted plasticity and responsiveness in MBON-α3. Importantly, while sensory input from Kenyon cells remained intact, concussions led to lasting dysfunction at the MBON level, suggesting that vulnerability to trauma arises within specific circuit nodes. These convergent findings provide a foundation for the discussion of how repeated head injury shapes brain function, plasticity, and long-term outcomes.

### Concluding remarks

Altogether, our study establishes a reproducible *Drosophila* model of repeated head trauma that reveals both transient and persistent consequences across behavior, cellular responses, and neural circuit function. We show that while flies can recover certain functions, such as locomotor capacity and MBON-α3 plasticity, other outcomes—including shortened lifespan, altered motivational state, glial proliferation, sleep architecture, and diminished MBON responsiveness—persist long after injury. These findings highlight the dual nature of recovery following concussions: the brain retains a capacity for functional plasticity yet also exhibits vulnerabilities that result in irreversible changes. By bridging behavioral phenotypes with cellular and circuit-level alterations, this work provides an experimental framework to dissect the mechanisms by which repeated concussions reshape neural systems and to explore potential strategies for mitigating their long-term impact.

## Materials and Methods

### Drosophila husbandry

Flies were cultured on standard medium at room temperature. Crosses, unless otherwise stated, were kept at 25 °C and 70% relative humidity with a 12-hr light-dark cycle. The following lines were used for experiments, crosses, and to generate stocks: *Canton-S*, *MCFO* (BDSC #64086; Nern et al., 2015), *r86e01-gal4* (Pfeiffer et al., 2011; Kremer et al., 2017), *MB082C-gal4* (Aso et al., 2014a), *uas-gcamp6f* (BDSC #42747, Chen et al., 2013); *uas-tdtomato* (BDSC #32221, Pfeiffer et al., 2011), *ok107*-*gal4* (BDSC #854; Connolly et al., 1996); and *uas-grabACh* (BDSC_86549; Jing et al., 2020).

### CTE protocol

*Drosophila* were loaded in a custom-built traumatic brain injury apparatus as described previously (Saikumar et al., 2020). Briefly, CO_2_-anesthetized flies were placed into a modified Heisenbeg collar to immobilize the flies by the head. A piezoelectric actuator (piezo.com #q220-a4-203yb) was used to deliver a compressive and precise injury to the fly head. We define three thresholds of injury as reported by Saikumar et al., (2020) –mild, moderate, and severe–, caused by compressing the fly head to 35%,40%, and 45% respectively. We included an additional sham group where flies were placed in the collar, but no head compression was applied. The repeated head compression protocol consisted of five repeated concussions with a one-minute ISI. After concussion, flies were removed from collar gently using a paintbrush and placed in food vials.

### Behavior Analyses

#### Survival curves

Flies were maintained in fresh food by flipping every two days. Every 24 hr at the same time (2 pm) vials were monitored to censor dead flies.

#### Climbing assay

Group of ∼20 concussed flies were loaded into a 20 mm ID-diameter Plexiglas cylinder. Flies were gently tap and their climbing behavior was recorded using a raspberry camera. The fraction of flies crossing a 50 mm mark by 18 s was registered. The assay was repeated 1, 5 and 10-days post-concussion.

#### Forced Swim Test

FST behaviors were evaluated by placing one fly per well in a 12-well culture dish containing 2 mL of 0.08% SDS at room temperature. The plate was illuminated from bellow using an infrared LED bed. An overhead FLIR camera containing an IR 720nm filter was used to record FST behaviors during the first 5 minutes in the well. Videos were then analyzed to quantify latency to first immobility, time spent immobile, and number of bouts of immobility. To verify that the flies were capable of mobility after the assay terminated, each fly was gently removed with a spatula and placed onto a paper towel. Any flies unable to immediately walk away from their landing locations were assumed drowned and subsequently excluded from final analysis.

#### Sleep

*Drosophila* sleep and activity were measured using the *Drosophila* activity monitoring system (Trikinetics) as described previously (Hendricks et al., 2000; PJ et al., 2000). In summary, flies were placed into individual 65 mm tubes and activity was continuously monitored. Locomotor activity was measured in 1 min bins and sleep was defined as period of quiescence lasting at least 5 min. Sleep in min/hr was plotted as a function of circadian time (hr) (Vecsey et al., 2024).

### Immunostaining

Whole brains were isolated and processed with minor modifications of those described (Jenett et al., 2012). Brains were first incubated with primary antibodies including: rabbit polyclonal anti-GFP (1:1,000, Life Technologies, cat# A11122), mouse monoclonal anti-nc82 (1:50, University of Iowa, DSHB), mouse monoclonal anti-FLAG (1:50, University of Iowa, DSHB), rat polyclonal anti-HA(1:800, Bethyl, cat# A190-118A) and chicken anti-V5 (1:1000, Roche, cat# 11867423001). Secondary antibodies included: anti-rabbit IgG conjugated to Alexa Fluor 488 (1:800, Life Technologies Cat# A11008, RRID: AB_143165), anti-mouse IgG conjugated to Alexa Fluor 633 (1:1000, Life Technologies Cat# A21052, RRID: AB_141459). anti-chicken IgY conjugated to Alexa Fluor 488 (1:1000, Invitrogen Cat# A32931), and anti-rat IgG conjugated to CFF633 (1:1000, Biotum, cat# 20137). Images were collected using a 20X objective with a Leica TCS SP8 confocal microscope with 488 and 633 nm laser excitation.

### In vivo imaging

Functional imaging was performed as previously reported (Martinez-Cervantes et al., 2022; Shah and Cervantes-Sandoval, 2025). Briefly, a singly fly was gently aspirated without anesthesia into a metal pipette connected to the vacuum (MicroGroup Hypodermic Tubing 304H22) to immobilize the head using proboscis aspiration. Once the head is immobilized, using a micromanipulator, the fly was inserted in a narrow slot the width of their body in a custom-designed recording chamber. The head was then fixed by gluing the sides of the eyes and thorax to the chamber using melted myristic acid. After this the fly was released from the metal pipette and the proboscis is fixed with myristic acid to avoid brain movement during proboscis extension. Using a syringe needle (BD PrecisionGlide^TM^ Needle REF 305193), a small, square section of dorsal cuticle was removed from the head to allow optical access to the brain. Fresh insect Ringer’s solution (103 mM NaCl, 3 mM KCl, 5 mM HEPES, 1.5 mM CaCl_2_, MgCl_2_, 26 mM NaHCO_3_, 1 mM NaH_2_PO_4_, 10 mM trehalose, 7 mM sucrose, and 10 mM glucose [pH 7.2]) was perfused immediately across the brain to prevent desiccation and ensure the health of the fly. Then the fat bodies and trachea above the brain was removed. Using a 20x water-immersion objective and a Leica TCS SP8 II confocal microscope with a 488 nm solid state laser, we imaged MBONs for 2 min at 2 Hz, during which odor stimuli was delivered starting 30 seconds after imaging initiation. We used one HyD channel (510-550 nm) to detect GCaMP6f fluorescence and 600-785nm to detect Tdtom. Recordings were performed at 2 Hz.

For MBON-α3, fluorescence was acquired from a region of interest (ROI) drawn around the dendrites of the neurons of interest. ΔF/Fo was calculated using a custom MATLAB script. Baseline was calculated as the mean fluorescence across the 5 s before each odor presentation. This baseline was then used to calculate %Δ*F*/*F*o for the complete recording. Bar graphs represent distribution of %Δ*F*/*F*_o_ responses across the 1 s of odor presentation. Solid lines in fluorescence traces represent mean %Δ*F*/*F*_o_ ± standard error (SE) (shaded area) across the odor responses.

### Odor presentation

To deliver odors to flies under the microscope, a stream of air (100 ml/min) was diverted (via solenoids) from flowing through a clean 20 ml glass vial to instead flow through a 20 ml glass vial containing a 0.5 µl drop of pure odorant. This air stream was then serial diluted into a larger air stream (1500 ml/min) before traveling through Teflon tubing (∼2.5 mm diameter) to reach the fly. Both solenoids that control odor delivery and the Grass stimulator that delivers shocks were controlled by Arduino microcontroller (Arduino Uno) with custom-made programs (available upon request). The regular training protocol consisted in twelve 1 s odor presentations of 3-octanol (OCT– Sigma, cat#218405) followed by three 1 s presentations of 4-methylcyclohexanol (MCH–Sigma, cat#589-91-3) with a 12 s ISI.

### Quantification and Statical Analysis

Statistics were performed using Prism9.0-Graphpad. All tests were two-tailed and confidence levels were set at α=0.05. The figure legends present all statistical tests use for each experiment and the p values.

---

**Figure 4 – figure supplement 1.**
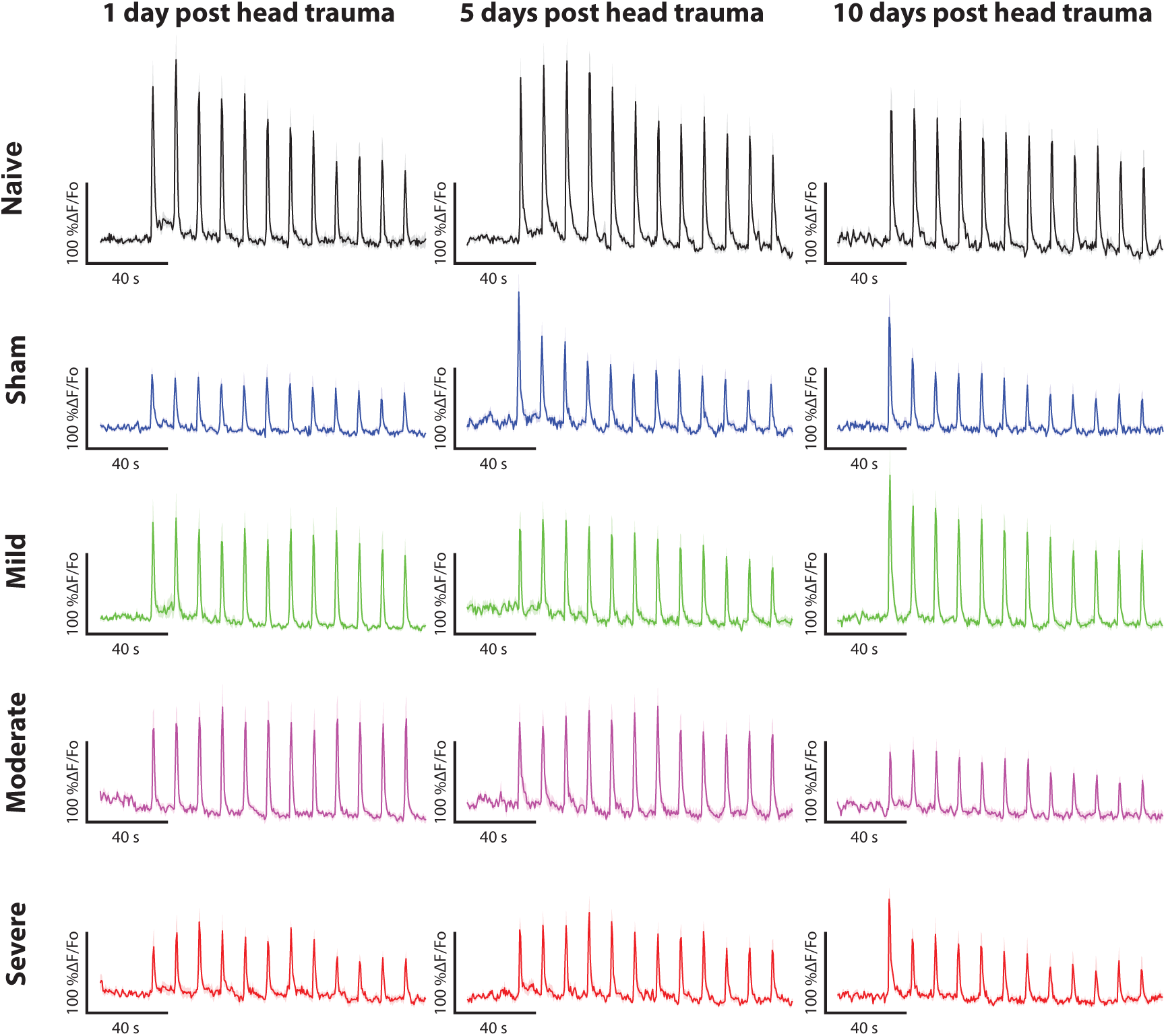
Raw olfactory responses to twelve 1 s odor presentations with a 12 s inter-stimulus interval in flies treated with sham, mild, moderate and severe repeated head trauma, one-, five-, and ten-days post-concussion. A age-matched naïve group is also showed.

